# QT-WEAVER: Correcting quartet distribution improves phylogenomic analyses despite gene tree estimation error

**DOI:** 10.1101/2024.11.11.622962

**Authors:** Navid Bin Hasan, Sohaib, Md. Shamsuzzoha Bayzid

**Author notes:** These authors contributed equally to this work (Navid Bin Hasan, Sohaib).

## Abstract

Summarizing individual gene trees into species phylogenies using coalescent-based methods has become a standard approach in phylogenomics. However, gene tree estimation error (GTEE) arising from a combination of reasons (ranging from analytical factors to more biological causes, as in short gene sequences) can potentially impact the accuracy of phylogenomic inference. We, for the first time, introduce the problem of correcting the quartet distribution induced by a set of estimated gene trees, which involves updating the weights of the quartets to better reflect their relative importance within the gene tree distribution. We present QT-WEAVER, the first method of its kind, which learns the conflicts within the quartet distribution induced by a given set of gene trees and generates an updated quartet distribution by adjusting the weights accordingly. QT-WEAVER is a general-purpose technique needing no explicit modeling of the subject system or reasons for GTEE or gene tree heterogeneity. Experimental studies on a collection of simulated and empirical data sets suggest that QT-WEAVER can effectively account for GTEE, which results in a substantial improvement in the species tree accuracy. Additionally, the concept of quartet conflicts and related algorithmic and combinatorial innovations introduced in this study will benefit various quartet-based computations. Therefore, QT-WEAVER advances the state-of-the-art in species tree estimation from gene trees in the face of GTEE. QT-WEAVER is freely available in open-source form at https://github.com/navidh86/QT-WEAVER.

## 1 Introduction

Species tree estimation from genes sampled across the whole genome has become routine with the advent of high-throughput sequencing technologies, generating genome-wide datasets that include hundreds or even thousands of loci. Species tree estimation from multiple genes is commonly done through concatenation (also known as “combined analysis”), which combines sequence alignments from different loci into a single supermatrix and then computes a tree on the super-matrix. While concatenation can produce accurate species trees when gene trees are concordant, it can be statistically inconsistent [9, 43], and produce incorrect trees with high support [24] when gene trees differ from the species tree due to various biological processes such as incomplete lineage sorting (ILS), gene duplication and loss (GDL), horizontal gene transfer (HGT), etc. As a result, summary methods [7, 20, 23, 26, 29–32, 36, 37, 50] that combine estimated gene trees while explicitly accounting for gene tree discordance have drawn substantial interest among systematists. However, summary methods are susceptible to gene tree estimation errors (GTEE), which can arise from various reasons including short gene sequences, inaccurate alignments, and the limitations of the models and algorithms used for inferring gene trees from sequence data.

Due to the growing awareness that GTEE is a major contributor to inaccuracies of summary methods, there has been great interest in developing tools [2, 8, 13, 17, 21, 25, 38–40, 51] to account for GTEE for improved species tree estimations. Most of these methods are species tree aware as they use a reference species tree in addition to the input gene trees. These species tree aware methods essentially reconcile/modify gene trees to make them closer in distance to the species tree by minimizing a species tree aware cost function. However, gene trees could be discordant and are not always expected to closely match the species tree. Additionally, “integrative” methods such as ProfileNJ [39] and TreeFix [51] use available sequence data as well as the estimated gene trees and reference species trees. Obtaining a reasonably accurate reference species tree despite substantial amounts of GTEE is difficult and the inaccuracy in the reference tree may have cascading effects on gene tree corrections. More importantly, species tree aware methods, specifically TreeFix and TRACTION [8], have been criticized for their potential to increase GTEE where gene trees are discordant with species trees [52]. Bayesian techniques for co-estimating both gene trees and species trees such as BEST [28], *BEAST [19], and PHYLDOG [6] can produce substantially more accurate gene and species trees than other methods, but are not scalable to genome-level analyses [4, 5, 27, 45, 49]. Therefore, despite significant attempts to account for GTEE, substantial challenges remain.

In this study, we address the problem of GTEE in the context of species tree estimation, by formulating the *Quartet Distribution Correction* (QDC) problem for the first time, where we seek to “correct/adjust” the distribution of quartets induced by a given set of estimated gene trees. QDC attempts to account for GTEE without resorting to any reference species tree. Quartet-based summary methods have drawn considerable attention because quartets (unrooted gene trees with four taxa) can avoid the “anomaly zone” [10–12], where the most likely gene tree topology may differ from the true species tree topology and can produce highly accurate species trees. ASTRAL, the most widely used summary method, infers a species tree by maximizing the number of quartets in the gene trees that are consistent with the species tree. Another class of methods (known as quartet amalgamation techniques [1, 32, 34, 41, 46]), such as wQFM [32] and wQMC [1], involves inferring weighted quartets for every group of four taxa (with weights correspond to the relative importance of the quartets) and then combining them into a single species tree. In this study, we present QT-WEAVER (**Q**uar**t**et **We**ight **A**djustment by **V**erifying, **E**stimating, and **R**ectifying quartet weights), a novel method that learns the weighted quartet distribution induced by a set of gene trees to identify and verify certain patterns of quartet conflicts and subsequently update the weights accordingly. We introduced the concept of “quartet conflicts” and proved key analytical and combinatorial results that underpin QT-WEAVER’s ability to learn and adjust quartet distributions without relying on any reference species tree or sequence data, offering a robust approach to improving the accuracy of species tree inference. The quartet distributions corrected by QT-WEAVER often align more closely with the true gene tree quartets than with those from the estimated gene trees, thereby effectively accounting for GTEE. Our experimental results, based on a diverse set of simulated and real biological datasets covering a wide range of challenging model conditions, demonstrate that amalgamating QT-WEAVER’s corrected quartets significantly improves species tree inference accuracy.

## 2 Quartet Distribution Correction Problem

### 2.1 Problem Definition

Let 𝒢= {*g*_1_, *g*_2_, …, *g*_*k*_} be a set of *k* gene trees, where each *g*_*i*_ is a tree on taxon set *S*_*i*_ ⊆*S* (i.e., any gene tree *g*_*i*_ can be on the full set *S* of *n* taxa or can be on a subset *S*_*i*_ of *n*_*i*_ taxa, making the gene tree incomplete). For a set of four taxa *a, b, c, d* ∈*S*, the quartet tree *ab* | *cd* denotes the unrooted quartet tree in which the pair *a, b* is separated from the pair *c, d* by an edge. Note that there are three alternative quartet topologies (*ab*|*cd, ac*|*bd, ad*|*bc*) for four taxa. Let 𝒬_*i*_ be the set present in 𝒢 be the set Therefore, 𝒬 = 𝒬_1_∪ 𝒬_2_ ∪ …∪ 𝒬_*k*_ is the multi-set of quartets present in 𝒢. Note that there are 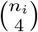 quartets in 𝒬_*i*_ and 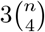 quartets in 𝒬 are unique, as there are three alternative quartet topologies for a set of four taxa. The gene tree frequency (GTF) based weighted quartet distribution 𝒬 𝒟 of 𝒢contains all possible quartets on *n* taxa along with their frequencies in the gene trees. That means, this is defined as the set 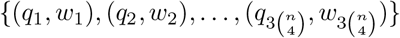 of 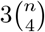 tuples.

Let 𝒢 _𝒯_ and 𝒢 _ℰ_ be the set of true and estimated gene trees, respectively. Consequently, the definitions of 𝒬_𝒯_, 𝒬_ℰ_, 𝒬 𝒟 _𝒯_, 𝒬 𝒟 _ℰ_ extend in an obvious way. We now define the quartet distribution correction (QDC) problem as follows.

#### Problem Quartet Distribution Correction (QDC)

Input A weighted quartet distribution 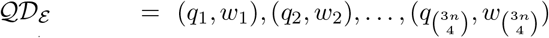 induced by a set 𝒢_ℰ_ of estimated gene trees.

Output A weighted quartet distribution 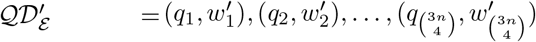 of updated weights 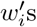 for *q*_*i*_s so that the divergence/difference between 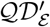 and the true distribution 𝒬 𝒟 _𝒯_ is minimized.

While we have defined the GTF-based weighted quartet distribution here, this concept extends to other weighting schemes as well.

### 2.2 Proposed Methodology: Identifying Quartet Conflicts

Our proposed approach is based on detecting patterns of “conflicts” among quartets, where a set of quartets is considered *conflicting* if they cannot be simultaneously satisfied by a single tree. Therefore, we try to learn the inherent patterns of conflicts within the quartets induced by the estimated gene trees and update the weights accordingly. We call the three alternative topologies (*ab*| *cd, ac*| *bd*, and *ad* |*bc*) for a set {*a, b, c, d*} of four taxa the *topological variants* of each other, and clearly, they are conflicting. We now formalize the concept of conflicts between quartets in Theorem 1, which generalizes how the number of common taxa between two quartets affects the number of trees that can simultaneously satisfy both quartets.

The leaves in a tree *T* are denoted by *L*(*T*). Every edge *e* in an unrooted leaf-labeled tree *T* defines a bipartition (or split) *π*(*e*) on the leaves *L*(*T*) (induced by the deletion of *e*).

#### Theorem 1

*Let q*_1_ *and q*_2_ *be two quartets drawn from a set of taxa. The number of binary trees on L*(*q*_1_) ∪ *L*(*q*_2_) *that simultaneously satisfy both q*_1_ *and q*_2_ *is determined by the number of taxa shared between them. Specifically, let ct denote the number of taxa common to both quartets, with ct* ∈{0, 1, 2, 3, 4}. *The number of binary trees that satisfy both q*_1_ *and q*_2_ *is as follows:*

#### Case 1

*ct* = 0, 1, 2 *When the number of common taxa between q*_1_ *and q*_2_ *is less than 3, i*.*e*., *ct* = 0, 1, 2, *there will always be multiple binary trees on L*(*q*_1_) ∪ *L*(*q*_2_) *that satisfy both quartets*.

#### Case 2

*ct* = 3 *When q*_1_ *and q*_2_ *share exactly three common taxa, meaning each quartet has one unique taxon relative to the other, two sub-cases arise:*

1. *If the unique taxon in q*_1_ *replaces the unique taxon in q*_2_ *(meaning that when the unique taxon in quartet q*_1_ *is replaced by the unique taxon in quartet q*_2_, *the resulting topology will match that of q*_2_*), there are exactly three distinct binary trees on L*(*q*_1_) ∪ *L*(*q*_2_) *that satisfy both quartets q*_1_ *and q*_2_.
2. *In all other configurations where the quartets share 3 common taxa, there is exactly one binary tree on L*(*q*_1_) ∪ *L*(*q*_2_) *that satisfies both quartets*.

#### Case 3

*ct* = 4 *When q*_1_ *and q*_2_ *share all 4 taxa (ct* = 4*), no binary tree can satisfy both quartets*.

*Proof*. We examine each case based on the number of common taxa between *q*_1_ and *q*_2_. Due to space constraints, we present the complete proof of this theorem in Appendix A1. However, we present the proof for Case 2 (*ct* = 3) here as our proposed algorithm for QT-WEAVER is based on this case (particularly Sub-case 2).

**– Case 2** *ct* = 3 **(three common taxa)** When *q*_1_ and *q*_2_ share exactly three taxa, the number of trees that can satisfy both quartets depends on the relationship between the unique taxon in each quartet. Two distinct sub-cases arise:

- Sub-case 1 (replacement of unique taxa): If the unique taxon in *q*_1_ simply replaces the unique taxon in *q*_2_ (preserving the relationship between the shared taxa) and vice versa, then two of the common taxa will be sister species in both *q*_1_ and *q*_2_. In this case, exactly three distinct binary trees on the set *L*(*q*_1_) ∪ *L*(*q*_2_) of five leaves satisfy both quartets. These three trees correspond to the possible ways to arrange the unique taxa from *q*_1_ and *q*_2_ with respect to the shared taxa (see Appendix A1 for detailed proof).
- Sub-case 2 (other configurations): In all other scenarios where the unique taxon in *q*_1_ does not simply replace the unique taxon in *q*_2_ (and vice versa), there is exactly one binary tree on the set *L*(*q*_1_) ∪ *L*(*q*_2_) of five taxa that satisfies both quartets. This is because the three common taxa, along with their relationships to the unique taxa, fully constrain the tree structure. Here, unlike sub-case 1, no pair of taxa among the three common taxa are sisters in both quartets. Consider the two quartets, *q*_1_ = *ab* |*cd* and *q*_2_ = *ae* |*bd* that satisfy this condition. In order to find a tree that satisfies both *q*_1_ and *q*_2_, we can insert *e* to *q*_1_ by making it a sister of *a*, as shown in Figure 1a. Similarly, we can insert *c* into *q*_2_ by making it the sister of *d* as shown in Figure 1b. Note that these two trees are identical (shown in shown in Figure 2a). Furthermore, the placements of *e* on four other edges in *q*_1_ (the edges incident on *b, c, d*, and the internal edge) result in trees that do not support *q*_2_ (*ae* |*bd*) as *b* becomes closer to *a* than *e* is to *a*. Similarly, the placements of *c* on four other edges in *q*_2_ do not produce any trees that satisfy *q*_1_ (*ab* |*cd*) as *b* becomes closer than *c* to *d*. Thus, there is exactly one tree (as shown in Figure 2a) that satisfies both *q*_1_ and *q*_2_.

**Fig. 1:**
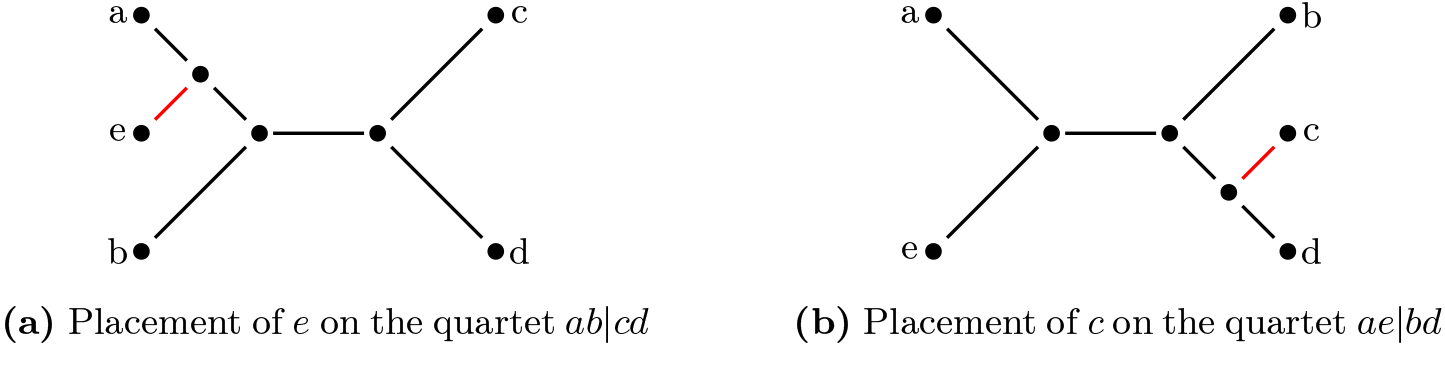
Unrooted trees satisfying both quartets *ab*|*cd* and *ae*|*bd*.

**Fig. 2:**
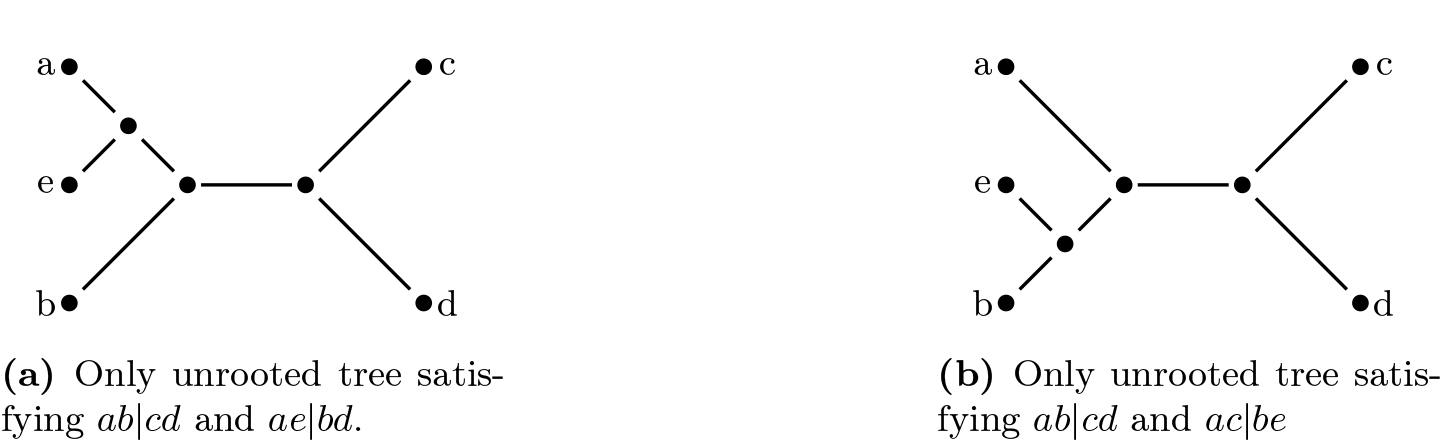
Unrooted trees satisfying canonical quartet pairs.

We define *canonical quartet pair* as a pair of quartets (*q*_1_, *q*_2_) that can be satisfied by exactly one tree *T*, where *L*(*T*) = *L*(*q*_1_) ∪ *L*(*q*_2_), as in Sub-case 2 of Case 2 in Theorem 1. Thus, a canonical quartet pair (*q*_1_, *q*_2_) uniquely represent a tree on *L*(*q*_1_) ∪ *L*(*q*_2_) that satisfy both *q*_1_ and *q*_2_. For example, *q*_1_ = *ab*|*cd* and tree *T* on {*a, b, c, d, e*} as shown in Figure 2a. There are 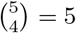 quartets (*ae*|*bd, q*_2_ = *ae*|*bd* is one such canonical quartet pair, which is satisfied by exactly one *ae*|*bc, ae*|*cd, ab*|*cd*, and *be*|*cd*) in *T* including the canonical pair corresponding to this tree (*ab*|*cd* and *ae*|*bd*). The canonical pair of quartets will clearly be in conflict with the topological variants of the three other quartets, *ae*|*bc, ae*|*cd*, and *be* |*cd*. Note that each quartet has two other topologically distinct variants. Thus, we have the following Corollary 1.

#### Corollary 1

*Every canonical pair of quartets* (*q*_1_, *q*_2_) *is in conflict with six quartets on L*(*q*_1_) ∪ *L*(*q*_2_).

A canonical pair and a quartet that is in conflict with this pair form a *conflicting set*. We now have the Corollaries 2-5. The proofs and related discussions are presented in Appendix A1.

#### Corollary 2

*Every binary unrooted tree T with five taxa contains four canonical pairs of quartets* (*q*_1_, *q*_2_), (*q*_2_, *q*_3_), (*q*_3_, *q*_4_) *and* (*q*_4_, *q*_1_) *with each of the four quartets q*_1_, *q*_2_, *q*_3_, *q*_4_ *being present in two canonical pairs*.

#### Corollary 3

*Every quartet q is part of eight canonical pairs of quartets, where the other member of each of these pairs has three common taxa and one unique taxon with respect to q*.

#### Corollary 4

*Every quartet q is part of 28 unique conflicting sets as a member of canonical pairs for each unique taxon x* ∈*/ L*(*q*).

#### Corollary 5

*For a set S of n taxa, every quartet q, where L*(*q*) ∈ *S, is in* 28 *×* (*n −* 4) *unique conflicting sets as a member of canonical pairs*.

Let {*q*_*i*_, *q*_*j*_, *q*_*k*_ } be a conflicting set, where *q*_*i*_ and *q*_*j*_ form the canonical pair in conflict with *q*_*k*_. Thus, while *q*_*i*_, *q*_*j*_, and *q*_*k*_ cannot all coexist in the same tree, any two of them can. Consequently, the presence of *q*_*i*_ in a species tree becomes less likely if the weights of both *q*_*j*_ and *q*_*k*_ increase (since higher weights for *q*_*j*_ and *q*_*k*_ indicate the possibility that both of them are present in the species tree). There are 28 · (*n −* 4) unique conflicting sets in each of which *q*_*i*_ is present (Corollary 5). By removing *q*_*i*_ from all these sets, we obtain a set of pairs *Z*_*i*_, where |*Z*_*i*_| = 28 · (*n −* 4).

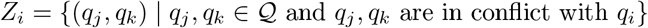

We now define the *conflict score* of a quartet as the sum of the products of the weights of pairs of quartets that are in conflict with it. Formally, for a given quartet *q*_*i*_ and a set *Z*_*i*_ containing the pairs of quartets that are in conflict with *q*_*i*_, the conflict score *C*(*q*_*i*_) is calculate follows:

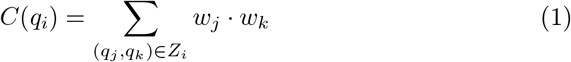

Note that in addition to the product of weights (PoW) based definition of the conflict score (Equation 1), we can define the conflict score using a minimum of weights (MoW) based approach as in Equation 2.

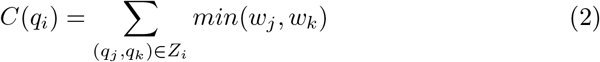

The rationale behind this MoW-based definition is that even if one of *w*_*j*_ or *w*_*k*_ is very large, it does not necessarily challenge the presence of *q*_*i*_ as long as the other is quite low. Therefore, considering the minimum of the two values (*w*_*j*_ and *w*_*k*_) as a contributor to the conflict score of *q*_*i*_ is a natural choice.

### 2.3 Algorithm: QT-WEAVER

Let 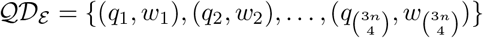 be the set of weighted quartets of a set 𝒢_*E*_ of estimated gene trees, *S* be the set of all taxa present in 𝒢_ℰ_ and 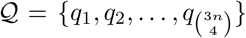 be the set of all quartets in 𝒬 𝒟_ℰ_. The algorithm takes the set ∈𝒬 𝒟_ℰ_ as input and for each quartet *q*_*i*_ ∈𝒬 𝒟_ℰ_, the algorithm adjusts the weight *w*_*i*_ to 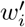 using the concept of quartet conflict we introduced in Section 2.2. We maintain a set 𝒜 ⊆𝒬, which represents the set of quartets whose weights have already been adjusted. Initially, 𝒜 =*∅*, indicating that no quartet weights have been adjusted at the start of the algorithm.

We iterate over the set of quartets, *q* ∈ 𝒬. If a quartet *q* is already adjusted, i.e., *q* ∈ 𝒜, we skip the quartet. Otherwise, let 𝒯= *q*_1_, *q*_2_, *q*_3_ be the set of three possible quartets on *L*(*q*). We then adjust the weights of the quartets in 𝒯 and insert them to, 𝒜 i.e., 𝒜 *←*𝒜 ∪ 𝒯 To adjust the weight for each quartet *q*_*i*_ in 𝒯 (*I* ∈{1, 2, 3}), we construct *Z*_*i*_, a set of all quartet pairs conflicting with *q*_*i*_. We do this by iterating over the set of taxa not present in *q*_*i*_, i.e. *S\ L*(*q*_*i*_). For each *t* ∈ *S \L*(*q*_*i*_), we obtain a set of 28 quartet pairs in conflict with the quartet *q*_*i*_ (Corollary 4). We add all such pairs to *Z*_*i*_. Thus, *Z*_*i*_ contains 28 *×*|*S\ L*(*q*_*i*_) | = 28 *×* (*n −*4) pairs of quartets. Next, we compute the *conflict score* for each of the quartets in 𝒯 using the PoW- or MoW-based formula in Equation (1) or Equation (2), respectively (this is a user defined configuration in our algorithm).

Note that the conflict score of a quartet *q*_*i*_ reflects its level of inconsistency relative to other quartets in the distribution. Thus, a quartet with a higher conflict score should be assigned a lower adjusted weight, reducing its relative importance due to its greater degree of conflict with others.

Therefore, we obtain 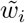 by dividing *w*_*i*_ by its conflict score *C*(*q*_*i*_).

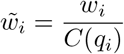

This operation on three quartets in 𝒯 denormalizes the weights of the quartets, which must be restored to their original sum *s* = *w*_1_ +*w*_2_ +*w*_3_. To normalize these weights, we multiply each by 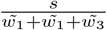 so that their sum equals their original sum *s*. The final normalized weight 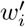 is given by:

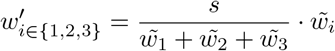

The pseudo-code for the QT-WEAVER algorithm is presented in Appendix A2

## 3 Experimental Study

### 3.1 Datasets

We evaluated the performance of QT-WEAVER using a collection of previously studied simulated and biological datasets. We studied two simulated datasets: a 37-taxon mammalian dataset based on biological data [47] and a 15-taxon dataset, both generated in prior studies [3, 35]. These datasets were created through a multi-stage simulation process, starting with a species tree, followed by the simulation of gene trees under the multi-species coalescent model (which can result in gene trees topologically distinct from the species tree), and finally, the simulation of gene sequence alignments down the gene trees. The datasets exhibit varying degrees of incomplete lineage sorting (ILS), ranging from moderate to high levels, and differ in the number of genes and the extent of gene tree estimation errors (controlled by sequence lengths). Thus, these simulated datasets provide a wide array of conditions under which we assessed the performance of QT-WEAVER. Table A3 in Appendix A4 presents a summary of these datasets. We also evaluated QT-WEAVER on a challenging biological avian dataset from Jarvis *et al*. [22] comprising 14,446 genes sampled from 48 birds.

### 3.2 Species tree estimation methods

We used wQFM, which was shown to be the most accurate quartet amalgamation technique [32, 33], to estimate species trees from weighted quartets. We also used wQMC, which is another well-known weighted quartet amalgamation technique. We ran GTF based wQFM and wQMC on uncorrected embedded quartets in the input gene trees with weights reflecting the frequencies of the quartets. wQFM (and wQMC) was also run on the adjusted/corrected weighted quartets, generated by QT-WEAVER, to demonstrate the impact of quartet weight correction on species tree estimation. We refer to this variant as *wQFM-corrected* or *wQFM+QT-WEAVER* interchangeably. We compared wQFM with ASTRALIII [36, 54] (v. 5.7.8), which is the leading quartet based species tree method. We also included the weighted variant of ASTRAL (wASTRAL [53]) in our study, which we configured to utilize weights derived from the branch support values in gene trees. The branch support values were estimated using non-parametric bootstrapping using RAxML [48]. Note that ASTRAL and wASTRAL cannot take a set of weighted quartets as input, and as such, we cannot evaluate these methods on corrected quartet distributions.

### 3.3 QT-WEAVER configurations

As discussed in Sections 2.2, 2.3, we can define the conflict score of a quartet *q* using PoW- or MoW-based approaches (Equations 1 and 2). Additionally, beyond using all 28 conflicting sets that a quartet *q* belongs to for each taxon *e* ∉ *L*(*q*), we explore subsets of these conflicting sets, analyzing subsets of sizes four, six, and eight. Thus, we have explored eight configurations (two weighting schemes combined with four conflicting set subsets). Our empirical exploration using the 15-taxon datasets (results presented in Appendix A3) suggests that the MoW-based approach is preferable over the PoW-based scheme. Moreover, utilizing six conflicting sets is both computationally faster and yields better performance across all conditions. Therefore, for the remaining experiments, we run QT-WEAVER with the configuration that uses 6 (out of 28) conflicting sets and the MoW-based approach to update quartet weights.

### 3.4 Measurements

We evaluated the accuracy of the estimated trees on simulated datasets by comparing them to the model species tree using the normalized Robinson-Foulds (RF) distance [42].Additionally, we assessed quartet scores, reflecting the number of quartets from the gene trees that are consistent with the estimated species tree. For the biological dataset, we compared the inferred species trees with those reported in the scientific literature. Multiple replicates under different model conditions were analyzed, and statistical significance between methods was determined using a two-sided Wilcoxon signed-rank test (with *α* = 0.05). To assess the accuracy of quartet distributions, we compared both corrected and uncorrected distributions to the true quartet distributions derived from the true gene trees using Jansen-Shannon divergence [16].

## 4 Results and Discussion

### 4.1 Results on 37-taxon dataset

The average RF rates of wQFM using both uncorrected and corrected (by QT-WEAVER) quartet distributions, ASTRAL, and wASTRAL on various model conditions in the 37-taxon dataset are shown in Figure 3(a). We vary the gene tree estimation error (by varying the sequence length from 50 to 1000 bp), the amount of ILS (by multiplying or dividing all internal branch lengths in the model species tree by two – producing three model conditions that are referred to as 1X (moderate ILS), 0.5X (high ILS) and 2X (low ILS), and the number of genes (from 100 to 500). In general, wQFM+QT-WEAVER is more accurate than wQFM across all model conditions – clearly demonstrating the benefit and positive impact of correcting quartet distributions for GTEE by QT-WEAVER. wQFM is, in general, more accurate than ASTRAL (which was also reported by prior studies [32, 33]). However, the improvements of wQFM over ASTRAL and wASTRAL are often not statistically significant. Remarkably, when wQFM is run on weighted quartet distributions corrected by QT-WEAVER, wQFM+QT-WEAVER becomes notably better than both ASTRAL and wASTRAL and in most cases (six out of nine), the improvements are statistically significant (*p* ≪ 0.05). Moreover, the accuracy achieved by ASTRAL and wQFM on true gene trees (i.e., without any estimation error) was matched by wQFM+QT-WEAVER even when using estimated gene trees derived from sequences as short as 500 bp. In fact, wQFM+QT-WEAVER with 1000 bp sequences is slightly more accurate than ASTRAL on true gene trees. These clearly show the power and efficacy of QT-WEAVER in accounting for gene tree estimation error. Note that, as the true gene trees do not have any corresponding gene sequences, non-parametric bootstrapping and subsequently wASTRAL is not applicable to this case. Even though there is no GTEE in true gene trees, the limited number of genes can result in a weighted quartet distribution that may fail to represent the true species trees. Interestingly, indeed, wQFM+QT-WEAVER outperformed both ASTRAL and wQFM on true gene trees, indicating that adjusting the weighted quartet distribution of true gene trees can still lead to more accurate species trees. We also evaluated the performance of wQMC, another well-known weighted quartet amalgamation technique, on quartets corrected by QT-WEAVER. Although wQMC is generally less accurate than wQFM [32, 33], we assessed wQMC+QT-WEAVER to demonstrate the usability of the corrected quartets generated by QT-WEAVER across different amalgamation methods. As shown in Figure A5 in Appendix A5.1, wQFM is better than wQMC (both for corrected and uncorrected data) and wQMC+QT-WEAVER consistently outperforms wQMC – further demonstrating the effectiveness of QT-WEAVER in adjusting quartet distributions and the superiority of wQFM over wQMC.

**Fig. 3:**
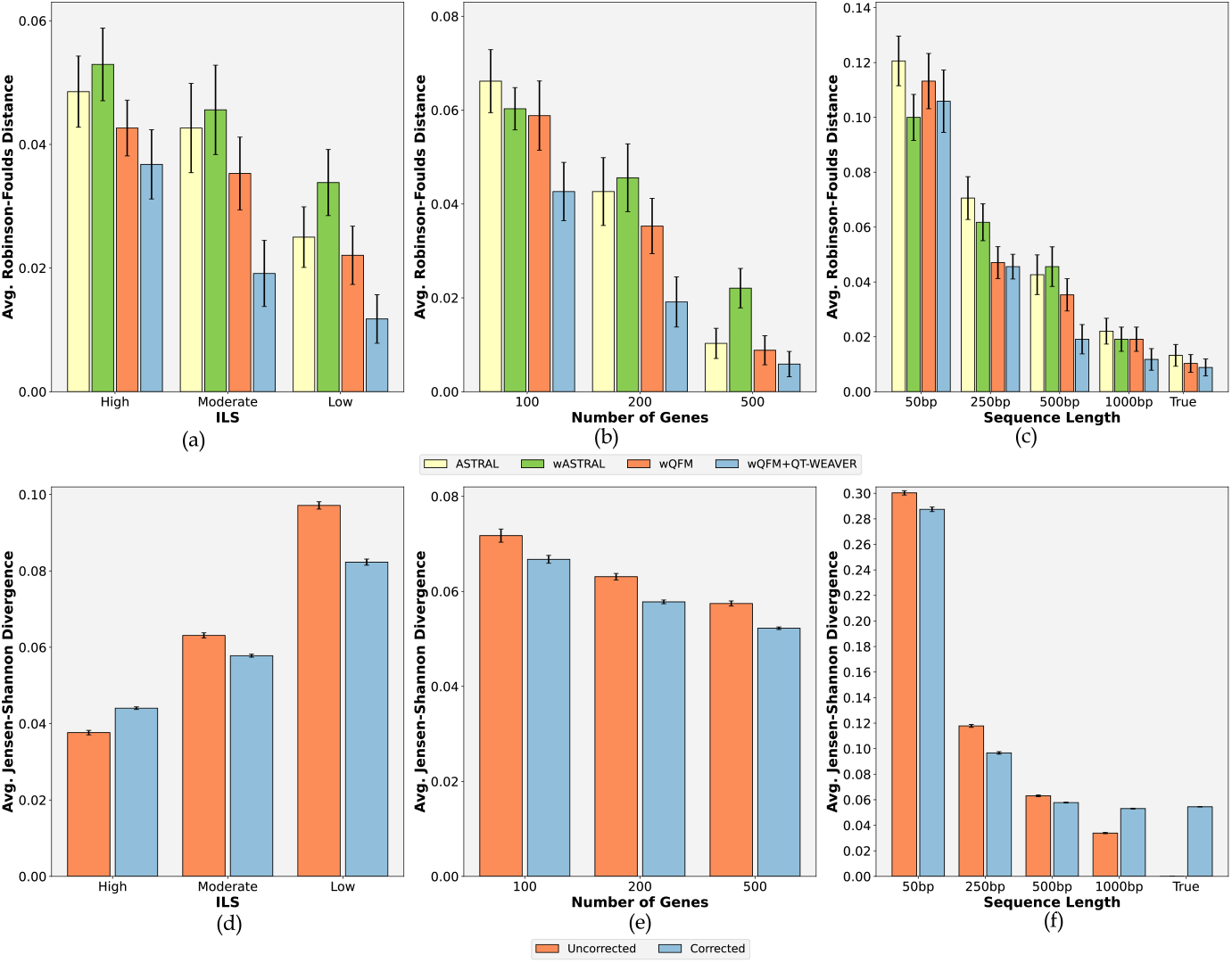
Results on 37-taxon dataset. (a)-(c) We show the average RF rates with standard errors over 20 replicates for the methods ASTRAL, wASTRAL (weighted variant of ASTRAL), wQFM (wQFM-uncorrected), and wQFM+QT-WEAVER (wQFM-corrected). (a) The amount of ILS was varied from high to low, keeping the number of genes fixed at 200 and sequence length at 500bp. (b) The number of genes was varied keeping ILS fixed at moderate (1X) and the sequence length at 500bp. (c) Varying sequence lengths with moderate ILS and 200 genes. We also include true gene trees with no GTEE. (d)-(f) Average Jensen-Shannon divergence of the quartet distributions before and after correction with respect to the true quartet distributions.

ASTRAL has been evolved and improved over successive versions for both accuracy and scalability, offering the theoretical guarantee of identifying the species tree that maximizes quartet support within the search space defined by the bipartitions of the input gene trees and thereby leaving limited (or no) scope for further enhancements in quartet score optimization. Thus, addressing GTEE becomes a critical avenue for pushing the accuracy frontier of quartet-based methods. In this context, the substantial improvements achieved by wQFM+QT-WEAVER over ASTRAL and wASTRAL is particularly noteworthy.

Next, we performed a series of experiments to further investigate the impact and quality of the quartet distributions produced by QT-WEAVER. First, to assess the similarity between the estimated quartet distributions (before and after correction) and the true quartet distributions (inferred from the true gene trees), we compare the Jensen-Shannon divergence between estimated and true quartet distributions in Figure 3(d)-(f) and Table A4 in Appendix A5.1. The corrected quartet distributions almost always have lower divergence than the uncorrected distributions, except for model conditions with very low divergence to begin with (less than 4 percent; 0.5x-200gt-500bp and 1X-200gt-1000bp conditions). The model conditions with relatively long sequence length (1000bp) have low amounts of GTEE and have less divergence to begin with (less than 4 percent), and correcting it does not seem to further reduce the divergence (rather making the distribution slightly more diverged). However, the corrected distributions overall result in better species trees as reflected by the RF rates (Figure 3(c)).

Overall, these results suggest that QT-WEAVER effectively bridges the gap with the overall nature of the true distribution.

Next, we computed the quartet scores of different estimated species trees and the true species tree with respect to both estimated and true gene trees (see Tables A5 and A6). When the quartet scores are computed based on original (uncorrected) estimated quartet distribution (Table A5), even though ASTRAL-estimated trees are less accurate than wQFM and wQFM+QT-WEAVER, ASTRAL achieves higher quartet scores than wQFM since ASTRAL is guaranteed to maximize the quartet score within a constrained search space. However, it “overshoots” the quartet score as it returns trees with higher quartet scores than the quartet score of the true tree. This is mostly because the statistical consistency of quartet score maximization criterion may not hold in the presence of gene tree estimation errors and limited number of genes. The true tree having the lower quartet scores across all model conditions supports this claim (see also [14, 15, 18] for more related discussions). The quartet scores of wQFM, especially wQFM+QT-WEAVER, with respect to the uncorrected estimated gene trees, are closer to the true quartet score than the scores of ASTRAL-estimated trees are to the true score. These further explain the superior performance of wQFM+QT-WEAVER over wQFM and ASTRAL. We also report the quartet score of wQFM+QT-WEAVER with respect to the quartet distribution corrected by QT-WEAVER. Notably, wQFM+QT-WEAVER has the highest quartet score when the score is computed based on the corrected quartet distribution. We note that these scores are not directly comparable to other methods’ quartet scores as other reported scores are computed with respect to a different quartet distribution (i.e., original uncorrected distribution). Therefore, we cannot consider these scores as “over-estimation” compared to true quartet scores. Rather, it suggests that the weight adjustment/correction by QT-WEAVER results in quartet distributions that may lead to species trees with higher quartet consistency scores (w.r.t. corrected distributions), which were not attainable with the uncorrected distributions.

Finally, we examined the quartet scores with respect to true gene trees with no estimation errors (Table A6). In most cases, wQFM and wQFM+QTWEAVER achieve higher and closer (to true quartet score) quartet scores than ASTRAL. This implies that the corrected distributions result in species trees that better correspond to the true gene tree distribution – supporting the trends observed in RF rates (Figure 3(a)).

### 4.2 Results on 15-taxon dataset

In the simulated 15-taxon datasets, we evaluated the performance on varying gene tree estimation errors using 100bp and 1000bp sequence lengths and on varying numbers of gene trees (100 and 1000). Similar to the 37-taxon dataset, wQFM+QT-WEAVER consistently outperforms ASTRAL, wASTRAL, and wQFM across all four model conditions (Figure 4). In particular, wQFM+QT-WEAVER is significantly better (*p* ≪ 0.05) than ASTRAL on challenging model conditions with short sequences (100bp), i.e., high GTEE.

**Fig. 4:**
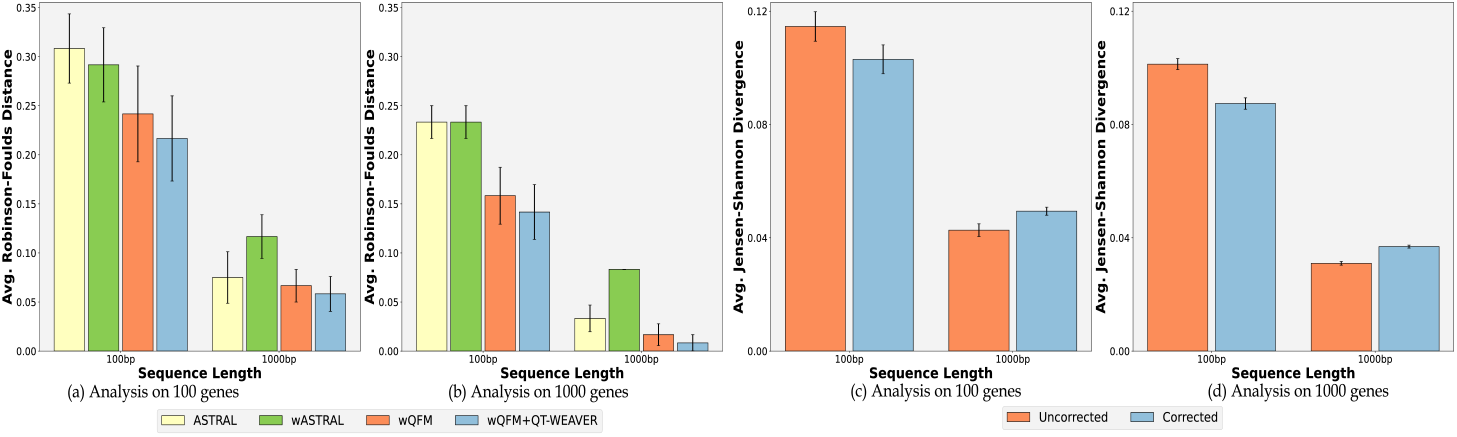
Results on 15-taxon dataset. (a)-(b) Average RF rates of ASTRAL, wASTRAL (weighted variant of ASTRAL), wQFM, and wQFM+QT-WEAVER on varying model conditions with varying sequence lengths and numbers of genes. (c)-(d) Average Jensen-Shannon divergence for the uncorrected and corrected quartet distributions.

Similar to 37-taxon dataset, we investigated the divergence of uncorrected and corrected quartet distributions from true quartet distributions (Figure 4(c)-(d) and Table A7) and the quartet scores of different estimated species trees with respect to both estimated and true gene trees (see Appendix A5.2).

## 5 Iterative corrections using QT-WEAVER

The demonstrated effectiveness of using QT-WEAVER in adjusting quartet distribution begs the question: what happens if we iteratively correct the adjusted distribution, using the output of QT-WEAVER as input to QT-WEAVER for the next iteration? To investigate this, we performed 50 iterations on 15-taxon dataset and analyzed how different evaluation metrics evolve in Figure 5. Notably, in model conditions with high gene tree estimation errors (100bp), RF rates steadily and dramatically improve (Figure 5a). For the 100bp-100gene condition, the RF rate drops from 25% to 11% after 15 iterations of weight corrections, and for the 100bp-1000gene condition, the RF rate decreases from 16% to as low as 2% after 10 iterations. Note that the RF rates of ASTRAL on these two model conditions are 31% and 23%, respectively, as indicated by the horizontal lines in Figure 5a. However, in model conditions with well-estimated gene trees (1000bp sequences) where the initial RF rates are already low (∼ 5%), repeated iterations may initially improve accuracy (e.g., a drop from 7% to 1% in the 1000bp-100gene condition) but eventually lead to performance degradation. This occurs because continuously adjusting well-estimated and corrected quartet distributions (as in cases with long sequences and large numbers of genes) can distort the weights to an extent that can mislead the tree search algorithm toward less accurate trees.

**Fig. 5:**
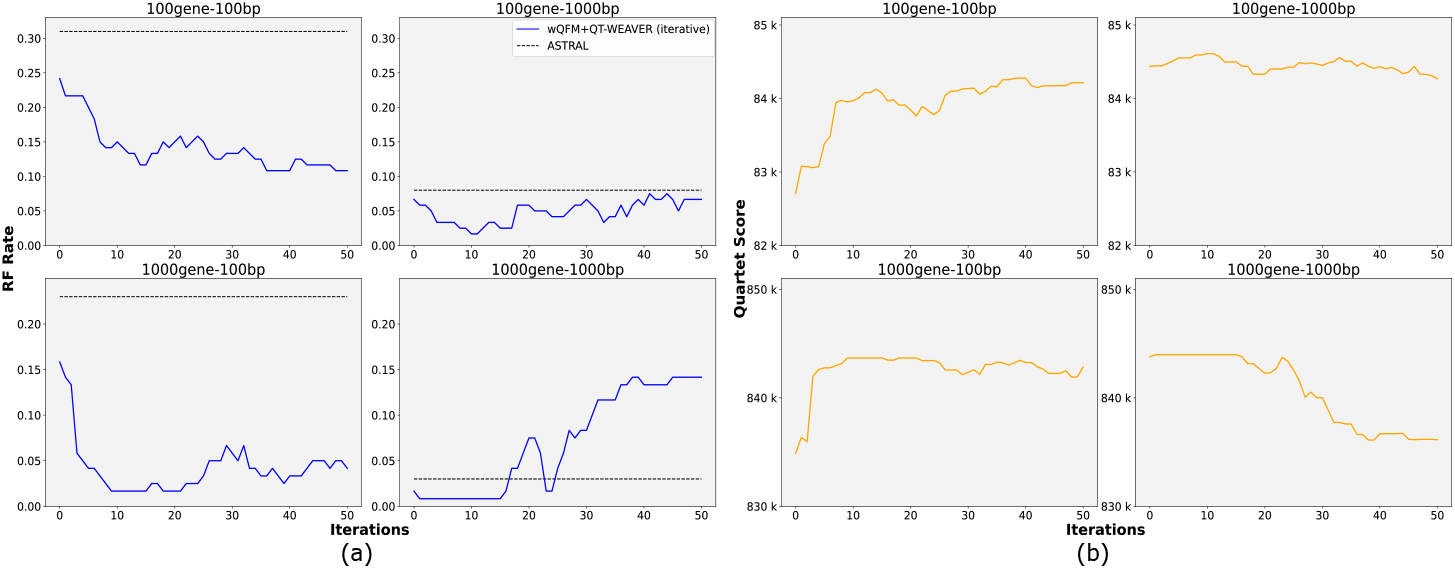
Iterative corrections on the 15-taxon dataset. We show the changes in (a) RF rates and (b) quartet scores (with respect to true gene trees) of wQFM+QT-WEVER over 50 iterations of weight corrections using QT-WEAVER. ASTRAL’s RF rates are indicated by dashed horizontal lines in (a).

In Figure 5(b), we show the changes in quartet scores (w.r.t. the true gene tree distributions) for wQFM+QT-WEAVER over multiple iterations. As we can see, the increase and decrease in the quartet scores over multiple iterations are perfectly aligned with the decrease and increase of the RF rates, respectively. When the GTEE is high, the scores maintain a rising trend. On the other hand, on the low GTEE model conditions, the quartet scores start to degrade after a few iterations in a pattern identical to the RF rate degradation. Additionally, we investigated the Jensen-Shanon divergence and quartet scores (relative to corrected distributions) across iterations (Figure A6 and related discussion in Appendix A5.3).

Thus, performing multiple iterations of weight adjustment shows potential, though improvements may not be consistent across all datasets. For instance, in the 37-taxon dataset, no further improvement was seen beyond the first iteration, as in most cases, the RF rates dropped below 5% after one iteration, and successive iterations distorted the distributions to an extent that resulted in worse trees. Nonetheless, QT-WEAVER’s iterative approach shows promise and warrants further investigation to develop a robust stopping criterion and scoring scheme for selecting the best tree from successive iterations.

## 6 Results on biological avian dataset

We have reanalyzed the avian biological dataset from [22], which has 14,446 genes (including exons, introns, and UCEs) across 48 taxa. This dataset contains high levels of gene tree discordance likely driven by rapid radiation events in the evolutionary history of these species. Mahbub et al. [32] compared ASTRAL, wQFM, and wQMC with the binned MP-EST tree which was presented in Jarvis *et al*. [22]. We include wQFM+QT-WEAVER in this comparison (Figure A7 in Appendix A6).

All three estimated trees are highly congruent with the reference binned tree, with wQFM+QT-WEAVER being the most congruent and ASTRAL the least.

The wQFM trees (on uncorrected and corrected distributions) are almost identical, with mainly one noticeable difference: wQFM+QT-WEAVER was able to correctly resolve the Cursores clade (crane and killdeer), which both wQFM and ASTRAL failed to recover. ASTRAL failed to recover Otidimorphae (bustard, turaco, and cuckoo), whereas both wQFM and wQFM+QT-WEAVER reconstructed this clade.

All three methods successfully reconstructed the well-established Australaves clade (passeriformes, parrots, falcons, and seriemas). They also recovered the Afroaves, Core Waterbird, and Caprimulgimorphae clades successfully. ASTRAL failed to recover Otidimorphae, unlike both the wQFM methods. All three failed to recover Columbea but were able to recover the constituent clades Columbimorphae (mesite, sandgrouse, and pigeon) and Phoenicopterimorphae (flamingo and grebe).

## 7 Running Time

The running time of QT-WEAVER solely depends on the number of taxa *n* and not on the number of genes, which determines the number of unique quartets 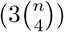 in the weighted quartet distribution. All analyses were run on the same machine with AMD Ryzen 7 5800H CPU (8 cores), 16GB RAM, and NVIDIA GeForce RTX 3060 GPU (6GB memory). The simulated 15-taxon and 37-taxon datasets took 1.65 and 242 seconds on average per replicate. The biological avian dataset with 48 taxa took 983 seconds to run.

## 8 Conclusions

This study, for the first time, introduces the quartet distribution correction problem and shows the impact and clear benefit of using quartet distributions corrected by QT-WEAVER for improved species tree estimations. QT-WEAVER learns the overall quartet distribution based on the pattern of quartet conflicts (a concept that we have introduced in this study), and seeks to update the weights to better reflect their relative importance. The concept of quartet conflict and related theoretical results have broad applicability and will be valuable for a range of quartet-based computational methods. Our experimental study shows that QT-WEAVER may result in substantial improvements over the leading method ASTRAL. Therefore, the idea of estimating species trees by correcting quartet distributions has merit and should be pursued and used in future phylogenomic studies. As a future study, we plan to evaluate QT-WEAVER on a diverse set of real biological datasets as the pattern of quartet conflicts are sufficiently complex and heterogeneous across various datasets. Another important research direction is the automatic selection of QT-WEAVER configurations. Additionally, considering the dramatic improvements achieved from multiple iterations of QT-WEAVER on certain datasets and model conditions, future studies need to investigate how to automatically identify an appropriate QT-WEAVER configuration for a given input, based on the input gene tree topologies, to ensure iterative enhancement.

## A1 Identifying Quartet Conflicts

### Theorem 1

*Let q*_1_ *and q*_2_ *be two quartets drawn from a set of taxa. The number of binary trees on L*(*q*_1_) ∪ *L*(*q*_2_) *that simultaneously satisfy both q*_1_ *and q*_2_ *is determined by the number of taxa shared between them. Specifically, let ct denote the number of taxa common to both quartets. Then the possible values of ct are 0, 1, 2, 3, or 4, and the number of binary trees that satisfy both q*_1_ *and q*_2_ *is as follows:*

### Case 2

*ct* = 3 *When q*_1_ *and q*_2_ *share exactly three common taxa (ct* = 3*), meaning each quartet has one unique taxon relative to the other, two sub-cases arise:*

1. *If the unique taxon in q*_1_ *replaces the unique taxon in q*_2_ *(meaning that when the unique taxon in quartet q*_1_ *is replaced by the unique taxon in the other quartet q*_2_, *the resulting topology will match that of q*_2_*), there are exactly three distinct binary trees on L*(*q*_1_) ∪ *L*(*q*_2_) *that satisfy both quartets q*_1_ *and q*_2_.
2. *In all other configurations where the quartets share 3 common taxa, there is exactly one binary tree on L*(*q*_1_) ∪ *L*(*q*_2_) *that satisfies both quartets*.

### Case 3

*Proof*. We examine each case based on the number of common taxa between *q*_1_ and *q*_2_.

**– Case 1** *ct* = 0, 1, 2 When *q*_1_ and *q*_2_ share fewer than 3 taxa, there are always multiple binary trees that satisfy both quartets.

1. Sub-case *ct* = 0 Since there are no shared taxa, the quartets are completely independent, and the quartets can be placed independently within a tree on *L*(*q*_1_) ∩ *L*(*q*_2_), leading to multiple distinct trees that satisfy both quartets.
2. Sub-case *ct* = 1 Suppose *q*_1_ = *ab* |*cd* and *q*_2_ = *ax* |*yz* be two quartets with *a* as the common taxon. Here, {a,b} and {a,x} are separated from {c,d} and {y,z} in *q*_1_ and *q*_2_, respectively. Then it is easy to see that any binary tree on *L*(*ab* | *cd*) ∪ *L*(*ax* |*yz*) = {*a, b, c, d, x, y, z*} containing an edge *e* that induces a bipartition *π*(*e*) = *abx* | *cdyz* separating {*a, b*}∪ {*a, x* }= {*a, b, x*} from {*c, d*} ∪{*y, z*} = {*c, d, y, z*} is consistent with both *ab*| *cd* and *ax*| *yz*. Since there are 3 possible ways to arrange {*a, b, x*} at one partition of *e* and 15 possible ways to arrange {*c, d, y, z* } in the other partition, there are at least 3× 15 = 45 trees that satisfy both quartets.
3. Sub-case *ct* = 2 Let the two common taxa in *q*_1_ and *q*_2_ be *a* and *b*, and the other four taxa (two from each quartet) be *c, d, x*, and *y*, respectively. Now, depending on the relative placements of *a* and *b* in *q*_1_ and *q*_2_, three cases may arise.
4. Sub-sub-case 1 (*a* and *b* are sister taxa in both quartets): Let *q*_1_ and *q*_2_ be *ab*| *cd* and *ab* |*xy* respectively. It is easy to see that any binary tree drawn on *L*(*q*_1_) ∪ *L*(*q*_2_) containing the bipartition *ab* |*cdxy* satisfies both quartets.
5. Sub-sub-case 2 (*a* and *b* are sisters in exactly one quartet): Without the loss of generality, let *q*_1_ and *q*_2_ be *ab* | *cd* and *ax* | *by, a* and *b* being sisters in *q*_1_. Then, any binary tree that contains both the bipartitions *ax* | *bcdy* and *abx* | *cdy* satisfies *ab* | *cd* and *ax* | *by*. Similarly, any binary tree that contains both the bipartitions *by* | *acdx* and *aby*| *cdx* satisfies *ab*| *cd* and *ax* | *by*.
6. Sub-sub-case 3 (*a* and *b* are not sisters in any quartet): Let *q*_1_ and *q*_2_ be *ac* |*bd* and *ax*| *by* respectively. Any binary tree on *L*(*q*_1_) ∪ *L*(*q*_2_) containing an edge *e* that induces a bipartition *π*(*e*) = *acx*| *bdy* separating {*a, c* } ∪ {*a, x*} = {*a, c, x* } from {*b, d*} ∪ {*b, y*} = {*b, d, y*} is consistent with both *ac*| *bd* and *ax* | *by*. Note that there are 9 such trees.

1. Sub-case 1 (replacement of unique taxa): If the unique taxon in *q*_1_ simply replaces the unique taxon in *q*_2_ (preserving the relationship between the shared taxa) and vice versa, then two of the common taxa will be sister species in both *q*_1_ and *q*_2_. In this case, exactly three distinct binary trees on the set *L*(*q*_1_) ∪ *L*(*q*_2_) of five leaves satisfy both quartets. These three trees correspond to the possible ways to arrange the unique taxa from *q*_1_ and *q*_2_ with respect to the shared taxa. For example, *q*_1_ : *ab*| *cd* and *q*_2_ : *ab*| *ce* satisfy this condition where *d* and *e* are the unique taxa in *q*_1_ and *q*_2_ respectively and replacing *d* with *e* in *q*_1_ (or *e* with *d* in *q*_2_) makes *q*_1_ and *q*_2_ identical. Moreover, *a* and *b* are closer to each other than they are to other species in both *q*_1_ and *q*_2_. In this case, we can place *e* on three branches in *q*_1_ (as shown in Figure A1a), resulting in three different trees on five taxa {*a, b, c, d, e*} that satisfy both of them. Similarly, we can place *d* on three branches in *q*_2_ (as shown in Figure A1b), producing three different trees that satisfy both *q*_1_ and *q*_2_. Note that the three trees in Figure A1a are identical to the trees depicted in Figure A1b. For instance, placing taxon *e* on the middle branch in Figure A1a yields the same tree as placing taxon *d* as the sister to taxon *c* in Figure A1b.

Let us now consider the two additional cases where taxon *e* is placed on the other two branches of the quartet *ab*| *cd*. First, if taxon *e* is placed as the sister taxon to *a*, one of the resulting induced quartets is *ae*| *bc*, which does not satisfy *ab*| *ce* as *e* is now the sister to *a*. Similarly, if *e* is placed as the sister to *b*, the quartet *ac*| *be* is formed, which also is not consistent with *ab*| *ce*. Thus, there are exactly three trees on {*a, b, c, d, e* } that satisfy both *q*_1_ and *q*_2_.

**Fig. A1:**
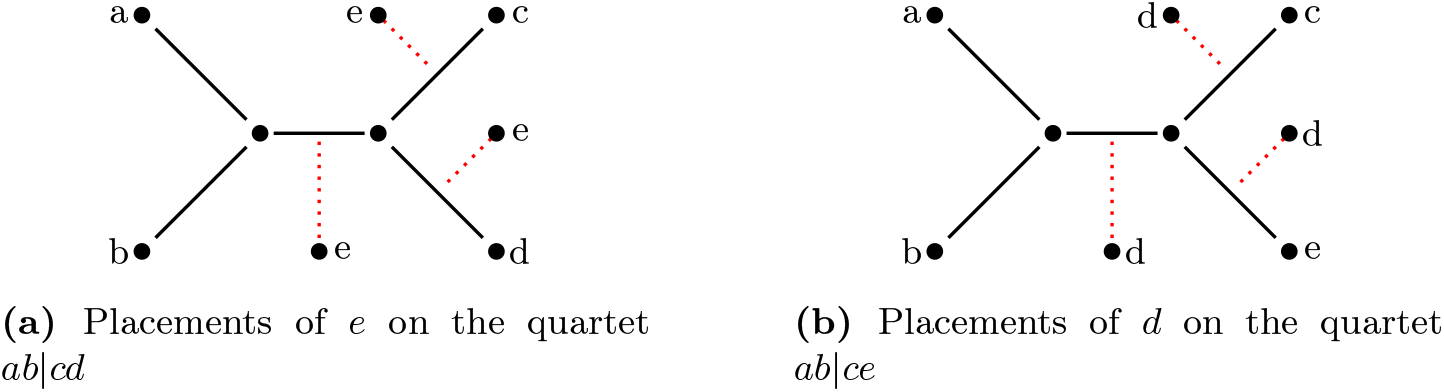
Placements for *e* and *d* on *q*_1_ and *q*_2_ respectively.

1. Sub-case 2 (other configurations): In all other scenarios where the unique taxon in *q*_1_ does not simply replace the unique taxon in *q*_2_ (and vice versa), there is exactly one binary tree on the set *L*(*q*_1_) ∪ *L*(*q*_2_) of five taxa that satisfies both quartets. This is because the three common taxa, along with their relationships to the unique taxa, fully constrain the tree structure. Here, unlike sub-case 1, no pair of taxa among the three common taxa are sisters in both quartets. Consider the two quartets, *q*_1_ = *ab* | *cd* and *q*_2_ = *ae* | *bd* that satisfy this condition. In this context, unique taxon *c* in *q*_1_ and *e* in *q*_2_ do not replace each other. In order to find a tree that satisfies both *q*_1_ and *q*_2_, we can insert *e* in *q*_1_ by making it a sister of *a*, as shown in Figure A2a. Similarly, we can insert *c* into *q*_2_ by making it the sister of *d* as shown in Figure A2b. Note that these two trees are identical. Furthermore, the placements of *e* on four other edges in *q*_1_ (the edges incident on *b, c, d*, and the internal edge) result in trees that do not support *q*_2_ (*ae* |*bd*) as *b* becomes closer to *a* than *e* is to *a*. Similarly, the placements of *c* on four other edges in *q*_2_ do not produce any trees that satisfy *q*_1_ (*ab* |*cd*) as *b* becomes closer than *c* to *d*. Thus, there is exactly one tree (as shown in Figure A3a) that satisfies both *q*_1_ and *q*_2_.

**Fig. A2:**
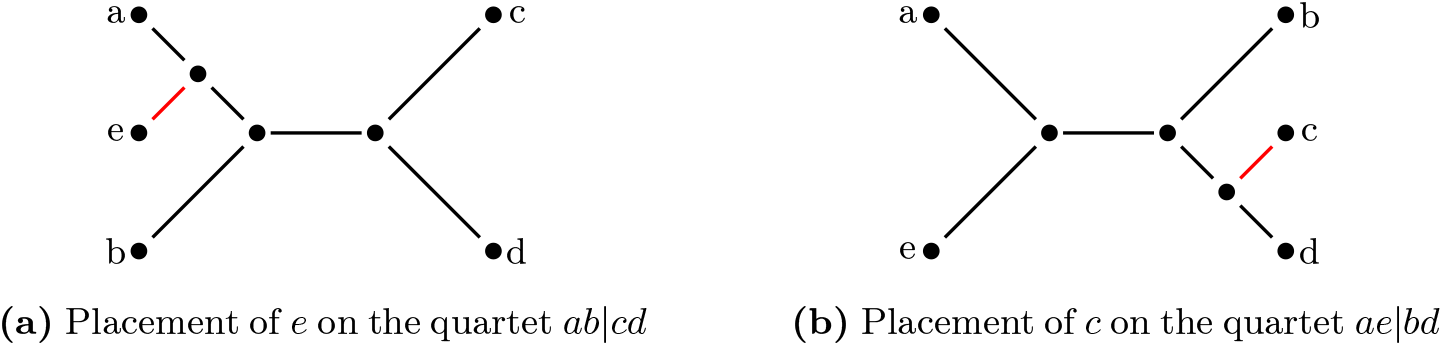
Unrooted trees satisfying both quartets *ab*|*cd* and *ae*|*bd*.

We define *canonical quartet pair* as a pair of quartets (*q*_1_, *q*_2_) that can be satisfied by exactly one tree *T*, where *L*(*T*) = *L*(*q*_1_) ∪ *L*(*q*_2_), as in Sub-case 2 of Case 2 in Theorem 1. Thus, a canonical quartet pair (*q*_1_, *q*_2_) uniquely represent a tree on *L*(*q*_1_) ∪ *L*(*q*_2_) that satisfy both *q*_1_ and *q*_2_. For example, *q*_1_ = *ab*|*cd* and tree *T* on {*a, b, c, d, e*} as shown in Figure A3a. There are *q*_2_ = *ae*|*bd* is one such canonical quartet pair, which is satisfied by exactly one *tree T on* {*a, b, c, d, e*} *as shown in Figure A3a. There are* 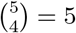 quartets (*ae*|*bd, ae*| *bc, ae* |*cd, ab*| *cd*, and *be*| *cd*) in *T* including the canonical pair corresponding to this tree (*ab* |*cd* and *ae*| *bd*). The canonical pair of quartets will clearly be in conflict with the topological variants of the three other quartets (*ae*| *bc, ae*|*cd*, and *be*|*cd*). Note that each quartet has two topologically distinct variants. Thus, we have the following Corollary 1.

**Fig. A3:**
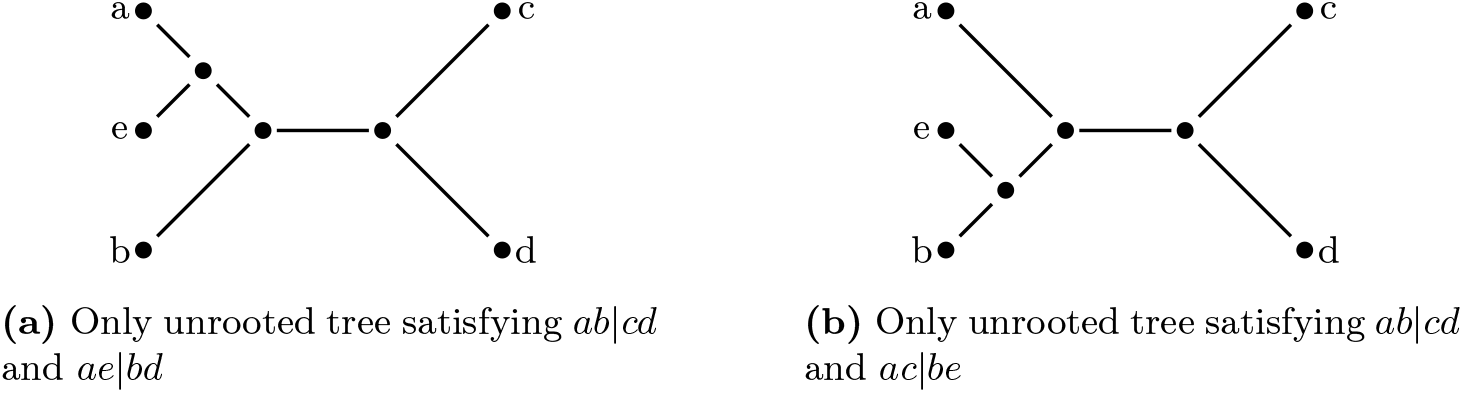
Unrooted trees satisfying two different canonical pairs.

### Corollary 1

Note that the tree in Figure A3a is constructed using *q*_1_ = *ab* |*cd* as the backbone, with the unique taxon *e* in *q*_2_ = *ae* |*bd* placed as a sister to *a*. Consequently, both *q*_2_ = *ae*| *bd* and *ae* | *bc* are induced by *T*. Therefore, if we instead consider *q*_2_ = *ae*| *bc* (rather than *ae*| *bd*), *q*_1_ = *ab*|*cd* and *q*_2_ = *ae* |*bc* forms another canonical pair, which is also satisfied by the same tree in Figure A3a. Thus, two canonical pairs, (*ab*| *cd, ae* |*bd*) and (*ab* |*cd, ae*| *bc*), that contain *ab* | *cd* as one member and the other member having *e* as the unique taxon with respect to *ab* |*cd* correspond to the tree shown in Figure A3a. Additionally, two other canonical pairs, (*ae*| *bc, be*| *cd*) and (*ae* |*bd, be*| *cd*), both sharing *be* |*cd* as a common member and excluding *ab*| *cd*, also represent the same tree in Figure A3a. Consequently, the tree shown in Figure A3a contains a total of four canonical pairs of quartets (*q*_1_, *q*_2_), (*q*_2_, *q*_3_), (*q*_3_, *q*_4_) and (*q*_4_, *q*_1_), where *q*_1_ = *ab*| *cd, q*_2_ = *ae* |*bd, q*_3_ = *be* |*cd, q*_4_ = *ae*| *bc*.

Notably, the quartet *ae* |*cd*, induced by the tree in Figure A3a, does not appear in any canonical pair, as there is no other quartet that can pair with *ae* | *cd* to place taxon *b* on the internal branch of *ae* |*cd*. This is because when two quartets form a canonical pair, the unique taxon from one quartet is positioned as a sibling of one of the taxa in the other quartet. All other quartets in the tree, except *ae* | *cd*, are part of two canonical pairs.

Additionally, using *q*_1_ = *ab* | *cd* as the backbone and placing *e* as the sister to *b*, we obtain a tree *T* (shown in Figure A3b) that satisfies two other canonical quartet pairs, (*ab*| *cd, ac* |*be*) and (*ab* |*cd, ad*| *be*), that contain *ab*|*cd* as one member. Similarly, *e* can be placed as a sister to the other taxa in *q*_1_ (i.e, *c* and *d*). Thus, for a given quartet *q*_1_ = *ab* |*cd*, we can form eight canonical pairs of quartets with *q*_1_ = *ab*| *cd* being one member and the other member *q*_2_ having *e* as the unique taxon to *q*_1_. These lead to the following Corollaries 2 and 3.

### Corollary 3

As stated in Corollary 1, a canonical quartet pair (*q*_1_, *q*_2_) is in conflict with six other quartets. Additionally, according to Corollary 3, *q*_1_ is a member of eight pairs of canonical quartets. Table A1 lists the six conflicting quartets for each of these eight canonical pairs of quartets, where *q*_1_ = *ab* |*cd* is one member and the other member contains a unique taxon *e*. Thus, every canonical pair and its six conflicting quartets form a set of three quartets that cannot coexist in a tree, resulting in 48 such sets. We call these conflicting trios of quartets as a *conflicting set*. Ignoring the duplicates, as marked in Table A1, 28 unique conflicting sets remain.

**Table A1:** List of conflicting quartets for eight canonical quartet pairs containing *ab* |*cd*. The items marked in blue are part of sets that were already encountered in a previous row. For example, the second quartet on the third row, *ae*| *bd*, constitutes the conflicting set {*ab*| *cd, ac*| *be, ae*| *bd*}. This set has already been encountered before. In the first row, by considering the second quartet *ac* |*be*, we obtain the conflicting set {*ab* |*cd, ae*| *bd, ac* |*be* }, which is identical to the set in consideration.

| Canonical Pair | Conflicting Quartets |
| --- | --- |
| $ab\vert cd$ & $ae\vert bd$ | $ab\vert ce$ \|\| $ac\vert be$ \|\| $ac\vert de$ \|\| $ad\vert ce$ \|\| $bc\vert de$ \|\| $bd\vert ce$ |
| $ab\vert cd$ & $ae\vert bc$ | $ab\vert de$ \|\| $ad\vert be$ \|\| $ac\vert de$ \|\| $ad\vert ce$ \|\| $bc\vert de$ \|\| $bd\vert ce$ |
| $ab\vert cd$ & $ac\vert be$ | $ab\vert de$ \|\| $ae\vert bd$ \|\| $ac\vert de$ \|\| $ad\vert ce$ \|\| $bc\vert de$ \|\| $bd\vert ce$ |
| $ab\vert cd$ & $ad\vert be$ | $ab\vert ce$ \|\| $ae\vert bc$ \|\| $ac\vert de$ \|\| $ad\vert ce$ \|\| $bc\vert de$ \|\| $bd\vert ce$ |
| $ab\vert cd$ & $ad\vert ce$ | $ac\vert be$ \|\| $ae\vert bc$ \|\| $ad\vert be$ \|\| $ae\vert bd$ \|\| $bc\vert de$ \|\| $be\vert cd$ |
| $ab\vert cd$ & $bd\vert ce$ | $ac\vert be$ \|\| $ae\vert bc$ \|\| $ad\vert be$ \|\| $ae\vert bd$ \|\| $ac\vert de$ \|\| $ae\vert cd$ |
| $ab\vert cd$ & $ac\vert de$ | $ac\vert be$ \|\| $ae\vert bc$ \|\| $ad\vert be$ \|\| $ae\vert bd$ \|\| $bd\vert ce$ \|\| $be\vert cd$ |
| $ab\vert cd$ & $bc\vert de$ | $ac\vert be$ \|\| $ae\vert bc$ \|\| $ad\vert be$ \|\| $ae\vert bd$ \|\| $ad\vert ce$ \|\| $ae\vert cd$ |

### Corollary 4

For a given set *S* of *n* taxa, and a quartet *q* (*L*(*q*) ∈ *S*), there are *n ™* 4 unique taxa in *S* with respect to *q*. Thus, the following corollary follows naturally.

## A2 QT-WEAVER: pseudo-code

### Algorithm 1 QT-WEAVER

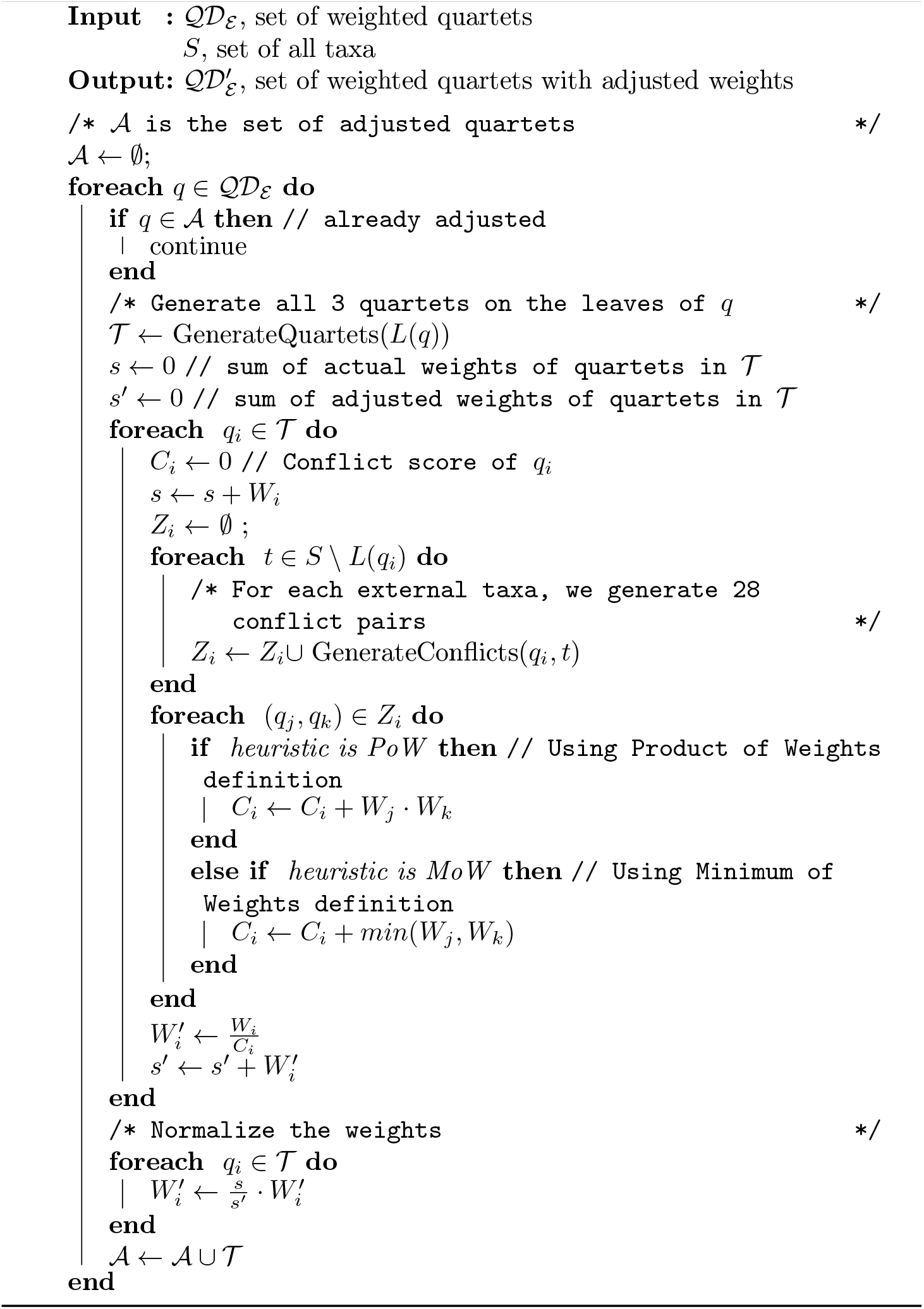

## A3 QT-WEAVER configurations

The comparison of the eight configurations of QT-WEAVER (the two weighting schemes combined with four different conflicting subsets) along with wQFM on the original distribution has been presented in Figure A4. The original distribution is derived by taking the frequency of each quartet in all gene trees (GTF: Gene Tree Frequency). We vary both the number of genes (100 and 1000) and the amount of gene tree estimation errors (by varying the sequence lengths from 100bp to 1000bp). As expected, the performance of the methods improves when the number of genes increases or when the gene tree estimation errors decrease. The MoW-based weighting scheme outperforms the PoW variant. Furthermore, subsets of four or six conflicting sets tend to perform better than all conflicting sets.

**Fig. A4:**
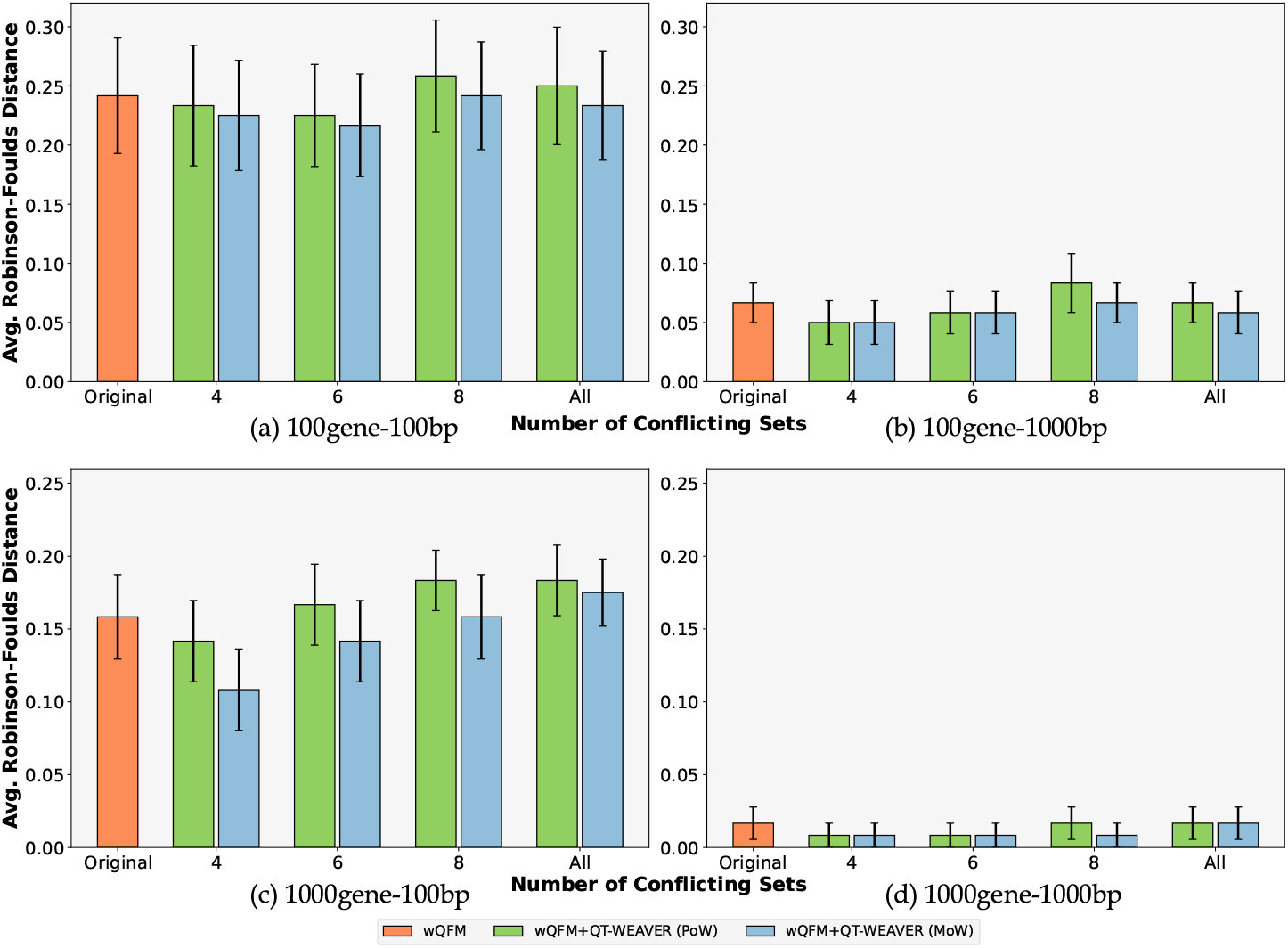
Results on 15-taxon dataset. Comparison of performance after correcting the weighted distributions using four, six, eight, and all 28 conflicting sets. We report the RF rates averaged over 10 replicates with standard errors. The distributions were amalgamated using wQFM. We show the effects of using both PoW and MoW weighting schemes.

These results raise the question – why does considering a small subset of all conflicting sets tend to yield better results? We hypothesize that using all 28 conflicting sets may impose excessive topological constraints on a quartet, making it challenging for QT-WEAVER to effectively distinguish among different topological variants.

In contrast, the six selected conflicting sets (as listed in Table A2) appear suitable for assessing conflict levels across quartet topologies, as indicated by our experimental results. While this specific choice of six sets lacks theoretical backing, we anticipate that the optimal subset–both in size and composition– may vary by dataset. Thus, an important research avenue involves automating the selection of these subsets, tailored to the topological features of the input gene trees, to optimize performance.

**Table A2:**
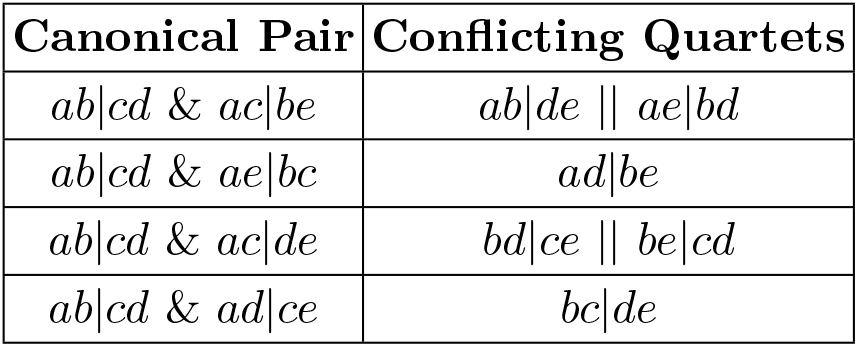
We list the six conflicting sets chosen from all the conflicting sets from Table A1.

| Canonical Pair | Conflicting Quartets |
| --- | --- |
| $ab\vert cd$ & $ac\vert be$ | $ab\vert de$ \|\| $ae\vert bd$ |
| $ab\vert cd$ & $ae\vert bc$ | $ad\vert be$ |
| $ab\vert cd$ & $ac\vert de$ | $bd\vert ce$ \|\| $be\vert cd$ |
| $ab\vert cd$ & $ad\vert ce$ | $bc\vert de$ |

## A4 Dataset

**Table A3:** Properties of the simulated datasets. The level of ILS is presented in terms of the average topological distance between true gene trees and true species trees.

| Dataset | ILS level | # genes | # sites | # Replicates | Ref. |
| --- | --- | --- | --- | --- | --- |
| 15-taxon | 82% | 100 - 1000 | 100 - 1000 | 10 | [3] |
| 37-taxon | 18%, 32%, 54% | 100 - 500 | 50 - 1000 | 20 | [35] |

## A5 Additional Results

### A5.1 Additional results on 37-taxon dataset

**Table A4:** The Jensen-Shannon divergence with respect to the true gene trees for different model conditions averaged over all replicates for the 37-taxon dataset. The best-performing method (i.e., the least amount of divergence) for each model condition has been highlighted in bold.

| Model Condition | GTF | QT-WEAVER |
| --- | --- | --- |
| 0.5X-200-500 | **0.038** | 0.044 |
| 1X-100-500 | 0.072 | **0.067** |
| 1X-200-25 | 0.354 | **0.345** |
| 1X-200-50 | 0.3 | **0.287** |
| 1X-200-250 | 0.118 | **0.097** |
| 1X-200-500 | 0.063 | **0.058** |
| 1X-200-1000 | **0.034** | 0.053 |
| 1X-500-500 | 0.057 | **0.052** |
| 2X-200-500 | 0.097 | **0.082** |

**Table A5:** Average quartet scores of estimated species trees and the true tree with respect to the estimated gene trees for the 37-taxon dataset. For wQFM+QT-WEAVER, we also report the quartet score with respect to the corrected estimated quartet distribution. The highest scores among the estimated trees have been italicized, and the scores closest to the true tree are shown in bold.

| Model Condition | ASTRAL | wQFM | wQFM+QT-WEAVER (Original Distribution) | wQFM+QT-WEAVER (Corrected Distribution) | True Tree |
| --- | --- | --- | --- | --- | --- |
| 0.5X-200-500 | 10007426 | 10006594 | **10006474** | *10415670* | 10003930 |
| 1X-100-500 | 5640132 | 5639605 | **5639168** | *5806039* | 5636884 |
| 1X-200-25 | 6491920 | 6489889 | **6489155** | *6662792* | 6481431 |
| 1X-200-50 | 7525465 | **7523379** | 7523903 | *7778394* | 7517956 |
| 1X-200-250 | 10569844 | 10568958 | **10568350** | *10941339* | 10565292 |
| 1X-200-500 | 11271938 | 11271426 | **11270680** | *11603816* | 11267991 |
| 1X-200-1000 | 11586641 | 11586355 | **11585945** | *11886743* | 11584970 |
| 1X-500-500 | 28171698 | **28171682** | 28168879 | *28983587* | 28171386 |
| 2X-200-500 | 11947266 | **11947020** | 11945419 | *12175146* | 11946371 |

**Table A6:**
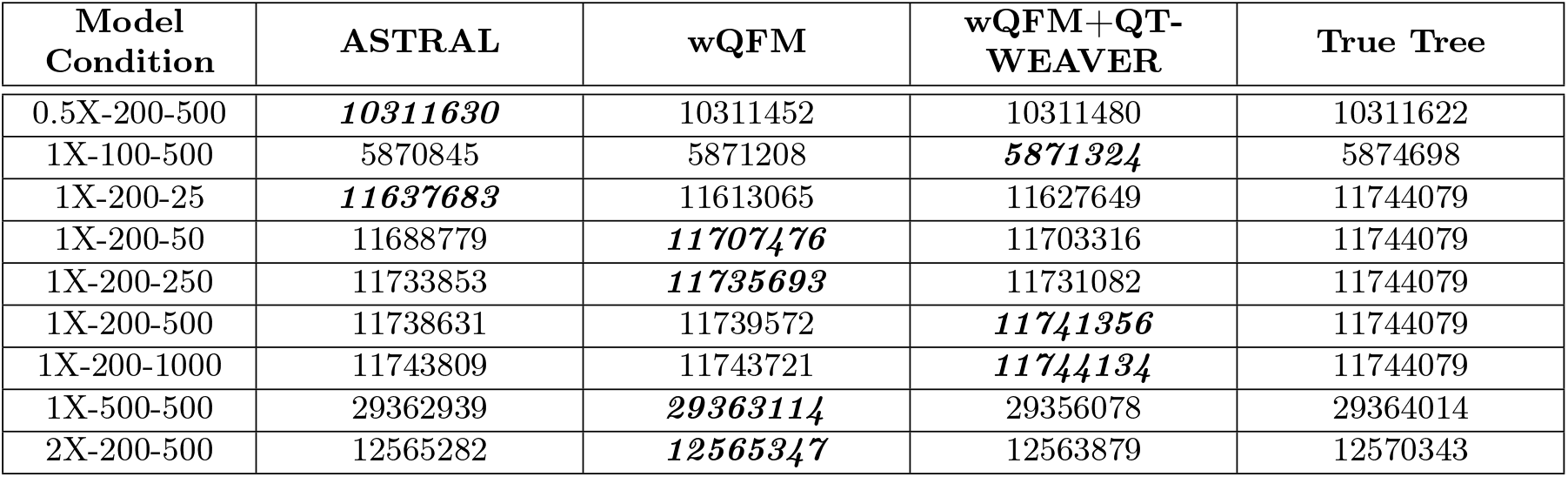
Average quartet scores of estimated species trees and the true tree with respect to the true gene trees for the 37-taxon dataset. The highest scores among the estimated trees have been italicized, and the scores closest to the true tree are shown in bold.

**Fig. A5:**
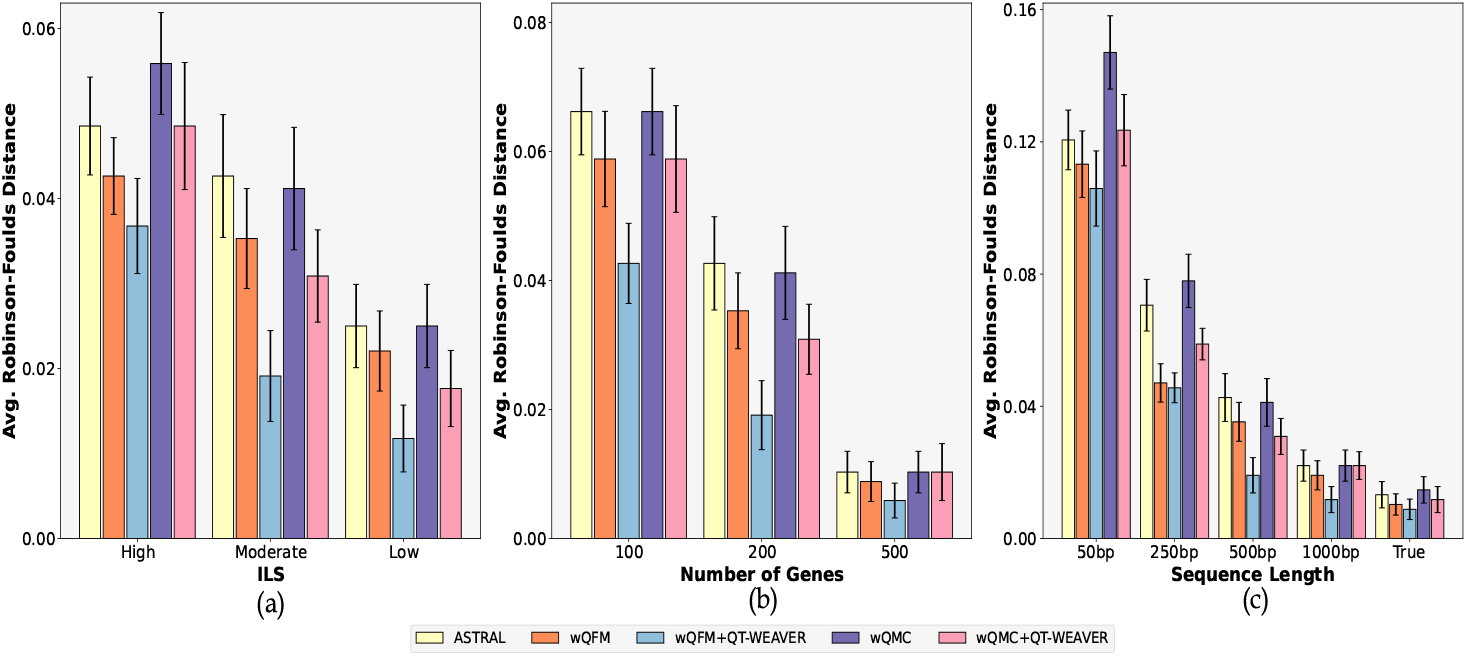
Results on 37-taxon dataset. We show the average RF rates with standard errors over 20 replicates for the methods ASTRAL, wQFM (wQFM-uncorrected), wQFM+QT-WEAVER (wQFM-corrected), wQMC (wQMC-uncorrected), and wQMC+QT-WEAVER (wQMC-corrected). The settings in (a)-(c) are identical to the ones in Figure 3.

### A5.2 Additional results on 15-taxon dataset

The Jensen-Shannon divergence clearly improves after correction when the gene tree estimation error is high (100bp). Thus, the quartet distributions inferred from gene trees with high estimation error realign with the true distribution after correction using QT-WEAVER. On the other hand, when the distribution has low amounts of GTEE with relatively long (1000bp) sequences and thus very low divergence to begin with (≤ 5%), correcting it does not seem to reduce the difference. Interestingly, even though the divergence increases slightly for the more accurate distributions (1000bp), the corrected distribution as a whole seems to be more representative of the true distribution, as supported by the better RF rates.

We again assess the quartet scores like we did for the 37-taxon dataset in Tables A8 and A9, and observed similar trends. wQFM+QT-WEAVER achieves higher and closer (to true quartet score) quartet scores than ASTRAL and wQFM across all four model conditions when the quartet scores are computed based on true gene trees – further demonstrating the efficacy of QT-WEAVER in accounting for GTEE.

**Table A7:** The Jensen-Shannon divergence with respect to the true gene trees for different model conditions averaged over all replicates for the 15-taxon dataset. The best-performing method (i.e., the least amount of divergence) for each model condition has been highlighted in bold.

| Model Condition | GTF | QT-WEAVER |
| --- | --- | --- |
| 100gene-100bp | 0.115 | **0.103** |
| 100gene-1000bp | **0.043** | 0.049 |
| 1000gene-100bp | 0.101 | **0.087** |
| 1000gene-1000bp | **0.031** | 0.037 |

**Table A8:** Average quartet scores of estimated species trees and the true tree with respect to the estimated gene trees for the 15-taxon dataset. For wQFM+QT-WEAVER, we also report the quartet score with respect to the corrected estimated quartet distribution. The highest scores among the estimated trees have been italicized, and the scores closest to the true tree are shown in bold.

| Model Condition | ASTRAL | wQFM | wQFM+QT-WEAVER (Original Distribution) | wQFM+QT-WEAVER (Corrected Distribution) | True Tree |
| --- | --- | --- | --- | --- | --- |
| 100gene-100bp | 69934 | **69777** | 69792 | *73598* | 69307 |
| 100gene-1000bp | 82167 | 82129 | **82125** | *87562* | 82099 |
| 1000gene-100bp | 693656 | 692949 | **692664** | *727712* | 690268 |
| 1000gene-1000bp | 818022 | 817994 | **817953** | *869904* | 817937 |

**Table A9:** Average quartet scores of estimated species trees and the true tree with respect to the true gene trees for the 15-taxon dataset. The highest scores among the estimated trees have been italicized, and the scores closest to the true tree are shown in bold.

| Model Condition | ASTRAL | wQFM | wQFM+QT-WEAVER | True Tree |
| --- | --- | --- | --- | --- |
| 100gene-100bp | 82159 | 82708 | ***83078*** | 84635 |
| 100gene-1000bp | 84345 | 84437 | ***84444*** | 84635 |
| 1000gene-100bp | 827970 | 834890 | ***836349*** | 844184 |
| 1000gene-1000bp | 843363 | 843792 | ***843973*** | 844184 |

### A5.3 Additional results on iterative corrections using QT-WEAVER

We investigated how the Jensen-Shannon divergence evolves across iterations (see Figure A6(a) in Appendix A5.3). Initially, the divergence decreases during the first one or two iterations but then begins to increase. This occurs because, for the three alternative quartet topologies (*ab* | *cd, ac* |*bd, ad*| *bc*) on four taxa, QT-WEAVER tends to prioritize the quartet topology it identifies as “correct”–the one with the least conflict score–by increasing its weight while reducing the weights of the alternative topologies. Thus, with successive iterations, the weight of the “correct” quartet topology continues to rise (toward 100%) while the weights of the other two alternatives keep decreasing (toward 0%). Thus, this overestimation of the weight of the “correct” quartets and the underestimation of the weights of the “incorrect” ones lead the adjusted distribution to diverge from the true weighted distribution. Despite this divergence, the iterative adjustments may still guide the tree search algorithm toward more accurate trees (up to a certain point) by emphasizing and amalgamating the quartets with higher weights, as evidenced by the gradual decrease in the RF rates up to 10-15 iterations. For the same reason, when examining the quartet scores of wQFM+QT-WEAVER with respect to the corresponding corrected quartet distribution (instead of the original quartet distribution), the scores continue to increase, approaching nearly 100% (see Figure A6(b) in Appendix A5.3). This occurs because, in the corrected distribution, the weights of the quartets present in the estimated species trees steadily rise, resulting in higher quartet scores for the estimated trees.

**Fig. A6:**
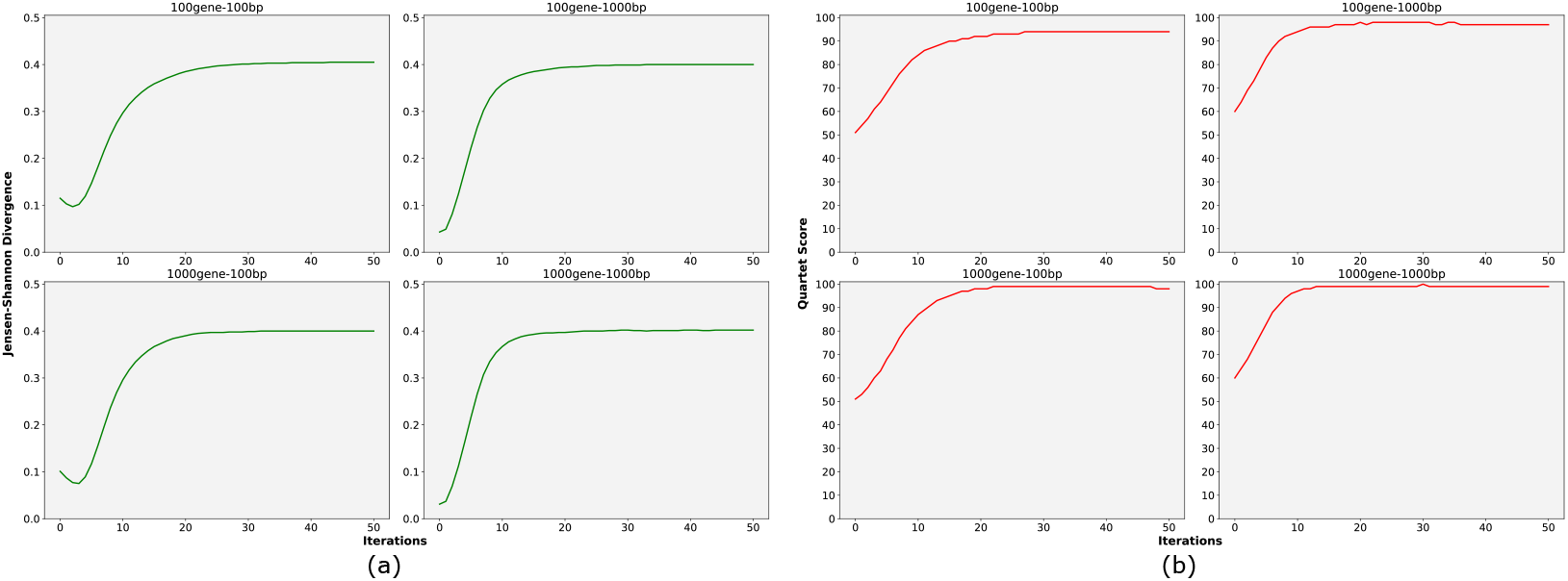
Results iterative corrections on 15-taxon dataset. We show the changes in (a) Jensen-Shannon divergence with respect to the true quartet distribution and (b) the quartet score of the estimated species tree with respect to its corresponding corrected quartet distribution over 50 iterations.

## A6 Analysis of avian biological dataset

**Fig. A7:**
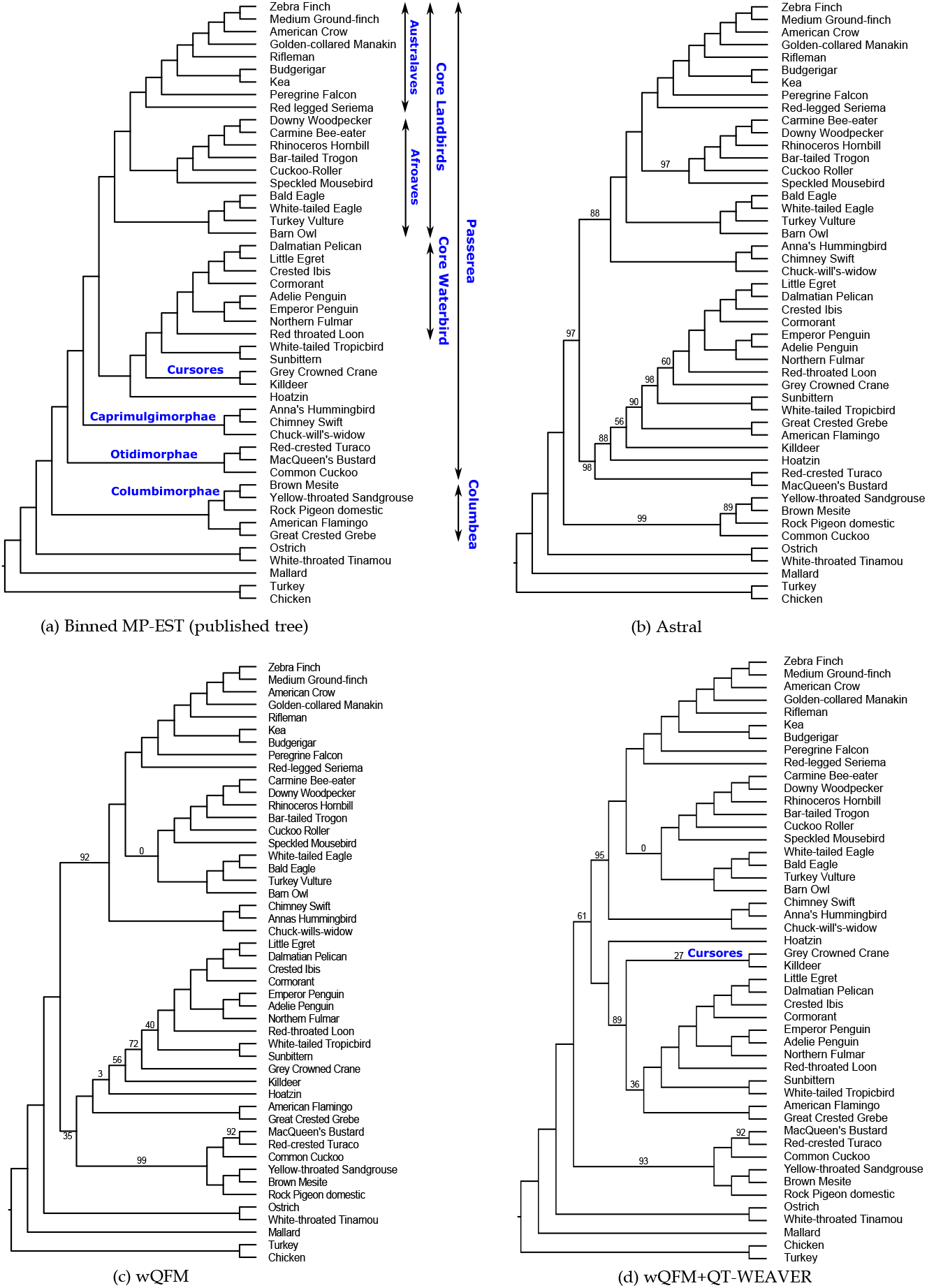
Analyses of the avian dataset using ASTRAL and wQFM (before and after correction). Branch supports are computed based on quartet-based local posterior probability [44] (multiplied by 100). All BS values are 100% except where noted

